# Functional amyloids in the microbiomes of a rat Parkinson’s disease model and wild-type rats

**DOI:** 10.1101/2021.03.31.438001

**Authors:** Line Friis Bakmann Christensen, Saeid Hadi Alijanvand, Michał Burdukiewicz, Florian-Alexander Herbst, Henrik Kjeldal, Morten Simonsen Dueholm, Daniel E. Otzen

## Abstract

Cross-seeding between amyloidogenic proteins in the gut is receiving increasing attention as a possible mechanism for initiation or acceleration of amyloid formation by aggregation-prone proteins such as αSN, which is central in the development of Parkinson’s disease. This is particularly pertinent in view of the growing number of functional (*i.e.* benign and useful) amyloid proteins discovered in bacteria. Here we identify two functional amyloid proteins, Pr12 and Pr17, in fecal matter from Parkinson’s disease transgenic rats and their wild type counterparts, based on their stability against dissolution by formic acid. Both proteins show robust aggregation into ThT-positive aggregates that contain higher-order β-sheets and have a fibrillar morphology, indicative of amyloid proteins. In addition, Pr17 aggregates formed *in vitro* showed significant resistance against formic acid, suggesting an ability to form highly stable amyloid. Treatment with proteinase K revealed a protected core of approx. 9 kDa. Neither Pr12 nor Pr17, however, affected αSN aggregation *in vitro*. Thus, amyloidogenicity does not *per se* lead to an ability to cross-seed fibrillation of αSN. Our results support the use of proteomics and formic acid to identify amyloid protein in complex mixtures and indicates the existence of numerous functional amyloid proteins in microbiomes.

**IMPORTANCE:** The bacterial microbiome in the gastrointestinal tract is increasingly seen as important for human health and disease. One area of particular interest is that of neurodegenerative diseases such as Parkinson’s which involve pathological aggregation into amyloid of human proteins such as α- synuclein (αSN). Bacteria are known to form benign or functional amyloid, some of which may initiate unwanted aggregation of *e.g.* αSN in the enteric nervous system through cross-seeding via contact with the microbiome. Here we show that the rat microbiome contains several proteins which form this type of amyloid aggregate both *in vivo* and *in vitro*. Although the two proteins we investigate in depth do not directly promote αSN aggregation, our work shows that the microbiome potentially harbors a significant number of bacterial amyloid which could play a role in human physiology at various levels.

## INTRODUCTION

Parkinson’s disease (PD) is a neurodegenerative movement disorder that affects 1-2% of all individuals above the age of 60 [1]. PD is characterized by the loss of dopaminergic neurons in the substantia nigra pars compacta (SNpc) of the midbrain and the accumulation of intracytoplasmic protein deposits referred to as Lewy bodies (LBs) or Lewy neurites (LNs) [2]. The LBs and LNs have been shown to be composed primarily of β-sheet-rich amyloid deposits of the intrinsically disordered protein α-synuclein (αSN) [3] which is believed to be a key player in the development of PD.

Within the cell, αSN is found in high concentrations at presynaptic structures [4, 5]. It has been suggested that αSN can spread via neuron-to-neuron propagation [6] and progression of sporadic PD has been proposed to happen as a caudo-rostral process (moving from the lower brain stem via the basal midbrain and forebrain to the cerebral cortex) [2]. Outside the central nervous system (CNS), LBs and LNs have also been observed in the enteric nervous system (ENS) in early stages of PD [7]. Recently it has even been proposed that there are two subtypes of PD, namely brain-first and body-first, depending on their place of origin [8]. Spreading from the ENS to the CNS has been suggested to happen through the vagus nerve as vagotomised individuals have a decreased risk for subsequently developing PD [9], and this has been confirmed in mouse models [10]. Further, enteric neurons secrete αSN under neuronal control [11]. Studies in rats have also shown that human αSN, injected into the intestinal wall, can be transported to the dorsal motor nucleus of the vagus via the vagal nerve [12]. Similarly, injection of recombinant αSN pre-formed fibrils into the gastric wall of mice was shown to result in LB pathology in the brainstem, and this was dependent on retrograde transport through the vagus nerve [13].

The great majority of neurons in the ENS is located in the myenteric and submucosal plexuses in the wall of the gastrointestinal (GI) tract [14]. Enteric nerve fibers extend through the different layers of the GI tract and have been shown to connect directly with enteroendocrine cells (EECs) [15]. Both EECs and intestinal epithelial cells express Toll-like receptors (TLRs) 1, 2 and 4 [16] and are therefore able to recognize different structures, often referred to as microbe-associated molecular patterns (MAMPs), expressed by, *e.g.,* gut bacteria. One example is the extracellular curli fibrils, which are aggregates of the CsgA protein produced by *Escherichia coli* and a wide range of other bacterial within the Enterobacteriaceae [17]. These functional amyloid fibrils are recognized by TLR1 and TLR2 and mediate interleukin 1β production [18]. This suggests that MAMPs produced by bacteria in our microbiome could interact with our CNS through the ENS, raising the possibility that αSN aggregation could be initiated through contacts with *e.g.* microbial proteins. In support of the pro-aggregatory role of CsgA, the amount of αSN deposits in aged rats and *C. elegans* PD models was increased after being orally fed with curli-producing *E.coli* compared to animals that had been fed with a curli-deficient *E. coli* strain [19]. Similar experiments have been performed in an transgenic (Tg) mouse model overexpressing αSN and again showed that exposure to curli correlated with increased αSN pathology in the gut and brain together with GI and motor deficits [20]. Introducing curli-producing *E. coli* in wild type (WT) mice had no effect on motor functions, suggesting that curli works in concert with other predisposing factors (like αSN overexpression) to cause disease [20].

Functional bacterial amyloid (FuBA) such as CsgA and its counterpart FapC in *Pseudomonas* species serve multiple roles in bacteria. They are unusually stable and often resist high concentrations of surfactant, denaturant or organic acids as well as being resistant to proteolysis. These robust properties may explain their ability to strengthen bacterial biofilms mechanically [21, 22] and increase their resistance to, *e.g.,* antibiotics [23]. Whether bacterial biofilms are actually formed in the (healthy) human GI tract is unclear. Conditions like the high flow rate through parts of the GI tract (2-4 h through the small intestine), gastric acid and the production of mucus by goblet cells are believed to make it difficult for bacteria to attach to at least the upper part of the digestive tract. However, the transit time is markedly longer in the colon (∼ 60 h) [24], and some bacteria produce hydrolytic enzymes that can break down the glycoproteins in the mucus and use the mucus layer as an energy and carbon source [25]. Many bacteria within the GI tract can form biofilm on medical devices like feeding tubes [26] and food particles [27]. *E. coli* strains isolated from the GI tract can produce both extracellular cellulose and curli [28].

Here we build on the observed correlation between FuBA and αSN aggregation. We analyse fecal samples from three transgenic PD rats and three WT rats for the presence of FuBA and their possible link to αSN aggregation. Due to the high stability of the FuBA structure, the amyloid state is insoluble under most denaturing conditions and only dissociate into monomers at high concentrations (80-100%) of formic acid (FA) [29, 30]. Thus, by treating complex samples with increasing concentrations of FA and identifying soluble proteins by trypsin digestion and LC-MS/MS analysis, potential FuBA can be identified by their markedly increased abundance at high (80-100%) FA concentrations [31]. In this way, we identify 365 candidates, which were further analyzed bioinformatically to narrow the list to the two most promising candidates (here called Pr12 and Pr17). These were expressed recombinantly and examined biophysically, demonstrating significant tendencies to form amyloid. However, Pr12/Pr17 seeds did not promote αSN fibrillation. Thus, amyloidogenicity *per se* does not imply an ability to cross-seed fibrillation of αSN.

## MATERIALS AND METHODS

### Materials

Fecal samples from three WT rats (WT1, WT2 and WT3) and three BAC-SNCA Tg Parkinson’s disease (PD) rats overexpressing human αSN (PD1, PD2 and PD3) [32] were kindly provided by Olaf Riess’s lab in Tübingen, Germany. All rats were female, three months old and kept in regular 12 h light/dark cycles. All animals had free access to both water and food.

### DNA extraction from WT and PD microbiome samples

Three WT samples and three PD samples were analyzed. DNA was extracted from the microbiome samples with a FastDNA spin kit (MP Biomedicals), following the manufacturers’ instructions. Briefly, 50 mg of fecal pellet was subjected to bead beating (4x40 s at 6 m/s with 5 min incubations in between) in a Lysing Matrix E tube (MP Biomedicals) using a FastPrep-24 instrument. Protein was precipitated, after which DNA was bound to a binding matrix suspension and eluted. DNA concentrations were determined with a Qubit dsDNA BR kit and the quality confirmed using gel electroporation on a 2200 TapeStation with D1000 Screentapes (Agilent Genomics).

### V1-V3 16S rRNA amplicon sequencing

All DNA samples were diluted to 5 ng/µL and mixed with barcode adaptors and a master mix containing dNTPs, MgSO4 and Platinum® Taq DNA polymerase high fidelity (Thermo Fischer Scientific) as previously described [33]. The following PCR conditions were used: 2 min incubation at 95°C followed by 30 cycles of [20 s at 95°C, 30 s at 56°C, 60 s at 72°C] and 5 min at 72°C. PCR products, now referred to as libraries, were purified using an Agencourt AMPure XP bead solution (Beckman Coulter) and a magnetic rack. Concentrations were determined with a Qubit dsDNA HS kit and amplification was confirmed on a 2200 TapeStation (Agilent Genomics). 30 ng of each library were pooled and submitted for MiSeq sequencing.

Forward reads were processed using usearch v.11.0.667. Raw fastq files were filtered for phiX sequences using -filter_phix, trimmed to 200 bp using -fastx_truncate -trunclen 200, and quality filtered using -fastq_filter with -fastq_maxee 1.0. The sequences were dereplicated using - fastx_uniques with -sizeout -relabel Uniq. Exact amplicon sequence variants (ASVs) were generated using -unoise3 [34]. ASV-tables were created by mapping the raw reads to the ASVs using -otutab with the -zotus and -strand both options. Taxonomy was assigned to ASVs using -sintax with -strand both and -sintax_cutoff 0.8 [35] and the Autotax processed SILVA 138 SSURef Nr99 database [36]. The raw sequencing data is available at the SRA with the accession IDs ERR3477180-ERR3477185.

### Direct identification of functional amyloid proteins in fecal samples by label-free quantitative MS

Functional amyloids were identified as previously described [31]. In short, 200 mg of each fecal pellet also used for DNA extraction were dissolved in 1 mL buffer (10 mM Tris-HCl, pH 8.0) containing 1X Halt™ protease inhibitor cocktail (Thermo Scientific). Cell lysis was achieved by bead beating in a Lysing Matrix E tube (MP Biomedicals) for 4x20 s at 6 m/s with 5 min breaks on ice in between beatings. Aliquots of 25 µL were transferred to 18 eppendorf tubes per sample (six different FA concentrations in three technical triplicates) before 1 h of lyophilization. Samples were then mixed with FA at concentrations of 0% (ultraclean water), 20%, 40%, 60%, 80% and 100% and lyophilized overnight. Lyophilized material was resuspended in a special reducing sodium dodecyl sulfate polyacrylamide gel electrophoresis (SDS-PAGE) loading buffer [37] and run on AnyKD gels (Biorad) for 10 min at 120 V, just allowing the samples to enter the gel. The gels were stained with Coomassie Brilliant Blue (CBB) G250. Gel bands were then cut into smaller pieces and reduced in a 10 mM DTT/0.1 M NH4HCO3 solution for 45 min at 56°C after several washing steps. In each washing step the gel pieces were rehydrated with 0.1 M NH4HCO3 for five minutes before addition of concentrated acetonitrile (1:1 v/v). Proteins were then alkylated for 30 min at room temperature (RT) in a solution of 55 mM iodoacetamide/0.1 M NH4HCO3. Finally, the gel pieces were washed thoroughly with 0.1 M NH4HCO3 and 1:1 v/v 0.1 M NH4HCO3/acetonitrile before drying in a vacuum centrifuge. In-gel digestion was performed overnight at 37°C by rehydrating the gel particles in 12.5 ng/µL trypsin in a 0.1 M NH4HCO3 buffer. Peptides were recovered mainly from the overnight buffer but also extracted from the gel particles by incubation in 5% FA followed by addition of acetonitrile. This extraction was performed twice and all supernatants were pooled and dried by vacuum centrifugation overnight. Peptides were reconstituted in 0.1% trifluoroacetic acid and 2% acetonitrile and subjected to ultra-performance liquid chromatography (UPLC) tandem mass spectrometry analysis by injection and concentration on a trapping column before separation on a separation column (Pepmap™ C18, Thermo Scientific) with a gradient of buffer B (100% acetonitrile) of 2% to 8% during the first minute and then from 8% to 30% in the following 39 min. Buffer B was then increased from 30% to 90% within 5 min. The UPLC system was coupled online to a Q Exactive Plus mass spectrometer (Thermo Scientific). The mass spectrometry data have been deposited to the ProteomeXchange Consortium (http://www.proteomexchange.org) via the PRIDE partner repository [38] with the dataset identifier PXD014649.

### Data analysis of LC-MS/MS data

Protein identification and quantification were performed as previously described [31] using the MaxQuant v1.5.8.3 software [39] and the label-free quantification (LFQ) algorithm [40]. The search was performed against the NCBI non-redundant protein sequence database. To prevent systematic errors, LFQ values were normalized between individual measurements and each protein is identified based on at least two detectable peptides. This allows for relative abundances to be compared across samples. Amyloid proteins are expected to give a sigmoidal appearance when their relative abundances are plotted against FA concentration (with higher FA concentrations, more protein monomer will be released from the fibrils and enter the SDS-PAGE gel). With an automated R- markdown script the data can be separated for each protein and the highest concentration will be given a value of 1. Hit proteins should fulfill the following requirements: 60 < f50 < 100, f’(f50) > 0.025 where f50 is the FA concentration required to depolymerize half of the amyloid fibrils and f’ (f50) is the slope of the fit at this concentration [31]. The fit used was: *f*(*x*) = 1⁄ (I)

### Bioinformatic analysis

Data analysis of the entire MS dataset were done using R [41]. All proteins were fitted as described [31]. We consider a protein to be a hit if at least one of the three triplicates shows a sigmoidal increase in abundance with FA concentration. Because well-characterized FuBA systems like Fap in *Pseudomonas* and curli in *E. coli* all require Sec-dependent secretion of the amyloid proteins, the hits were all analyzed by the SignalP 4.1 algorithm [42] to restrict the list to candidates containing a signal peptide. RADAR [43], AmylPred2 [44] and Clustal2.1 were used to identify imperfect repeats, amyloidogenic amino acid sequences and sequence alignments, respectively.

### Expression and purification of candidate amyloid proteins

pET30a vectors encoding proteins WP_032523104.1 and OAD22177.1 (referred to as Pr12 and Pr17, respectively) were prepared by Genscript. Both proteins were produced without signal peptides (residues 1-22 for Pr12 and residues 1-18 for Pr17) but with an N-terminal His6 tag. *E. coli* BL21(DE3) bacteria were transformed with the expression vectors, spread on LB agar plate with 50 µg/ml kanamycin, grown up 24 h and then transferred to LB medium containing 0.1% glycerol, 50 µg/ml kanamycin and 4 mM MgSO4. Cells were grown up in an incubator at 37°C and 125 rpm and induced with 1 mM IPTG at OD600 of 0.6 – 0.8. After 3 hrs, cells were harvested by centrifugation at 4000 g for 20 minutes. Pellets were suspended in 50 ml of denaturation buffer (8 M GdmCl, 50 mM Tris-HCl, pH 8.0) with one protease inhibitor tablet (Roche) per L medium, lysed by slow stirring of the GdmCl-cell pellet suspension overnight at 4°C and spun down at ∼ 12,500g for 30 min at 15°C. The supernatant was incubated with nickel-nitrilotriacetic acid beads (typically 45 ml supernatant to 5 ml beads) for 1 h at 4°C on rolling table with 60 rpm, after which the beads were packed in an empty PD10 column and washed with washing buffer (8 M GdmCl, 50 mM Tris-HCl, 30 mM imidazole, pH 8.0) before elution of the protein into elution buffer (8 M GdmCl, 50 mM Tris-HCl, 300 mM imidazole, pH 8.0). Protein elution fractions of ∼ 2 mL were immediately frozen in liquid nitrogen and stored at -80°C.

### Purification and preparation of αSN

αSN was prepared as described [45] and protein concentration was determined at 280 nm using an extinction coefficient of 5,960 M^−1^ cm^−1^ and the molecular weight of 14,460 Da after the lyophilized αSN had been dissolved in 60 mM Tris buffer, pH 7.4, and filtered through a 0.2 µm filter.

### Thioflavin T (ThT) assays of Pr12 and Pr17

The two proteins were first desalted into 1X PBS, pH 7.4, and protein concentration was determined at 280 nm using extinction coefficients of 14,100 and 14,690 L mol^-1^ cm^-1^ for Pr12 and Pr17, respectively, based on their amino acid composition. The proteins were then immediately transferred to a 96-well Nunc optical bottom plate (Corning) containing 40 µM ThT. Fibrillation was followed using 445 nm excitation and 485 nm emission at 37°C on an Infinite Pro 23 (Tecan, Männersdorf) plate reader. Due to precipitation issues in 1X PBS, we started desalting the proteins from the elution buffer (8 M GdmCl, 50 mM Tris-HCl, 300 mM imidazole, pH 8.0) into 4 M urea in 1X PBS or 60 mM Tris, pH 7.4 (Pr12) or 8 M urea in 1X PBS, pH 7.4 (Pr17) to keep the proteins monomeric. Using Amicon Ultra spin filters (Merck) with a 3 K (Pr12) or 10 K cutoff (Pr17) both proteins were concentrated to either 7.5 or 10 mg/mL. The aggregation behaviour of the two proteins (final concentration of 0.5 mg/mL) were then investigated in the presence of different urea concentrations or at different pH values. Because the proteins had been concentrated in urea, 0.2 M urea (Pr12) or 0.4 M urea (Pr17) were also present in all ThT assays. The buffers used to investigate pH effects were 80 mM citric acid (pH 3), 60 mM citric acid (pH 4), 60 mM MES (pH 6), 1X PBS (pH 7.4) and 60 mM Tris (pH 8).

### Size-exclusion chromatography (SEC) analysis of urea-treated Pr17

After fibrillation of Pr17 in the presence of increasing [urea], the samples were subjected to size-exclusion chromatography using a Superose 6 10/300 GL column (GE Healthcare). For each sample, the column was equilibrated with a buffer containing the same urea concentration in 1X PBS as the sample to be investigated. Then, 500 µL sample was injected and run with a flow rate of 0.5 mL/min on an ÄKTA Pure protein purification system (GE Healthcare). Control samples containing ThT alone at the lowest (0.4 M urea) and the highest (6 M urea) were also included.

### Stability studies of Pr12 and Pr17 aggregates

Pr12 and Pr17 fibrils were produced by desalting the protein directly into 60 mM MES, pH 6, and incubating the samples overnight at 37°C at 300 rpm. Aggregates were pelleted by centrifugation (13,500 rpm, 15 min) in a MicroStar12 table centrifuge (VWR) and washed twice in milli-Q water (MQ) before equal volumes of aggregates were aliquoted into new tubes, Pellets were then treated with equal amounts of solutions containing increasing [urea] or [FA] and incubated for 10 min at RT before pelleted again. 20 µL of the supernatant was then transferred to new tubes and either mixed with R loading buffer (the urea-treated samples) or subjected to lyophilization (the FA-treated samples). After lyophilization the samples were mixed with 20 µL of a special loading buffer containing 8 M urea. Both the urea- and FA-treated samples were analysed with SDS-PAGE and quantified with ImageJ (https://imagej.net/). The percentage of solubilized Pr12/Pr17 aggregates was normalized to the 100% FA band and fitted to the following equation:

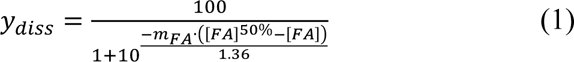

where [FA]^50%^ is the FA concentration where half of the aggregates are dissolved, i.e. the two states (aggregated state and monomeric state) are equally stable. The *m*-value, as in conventional protein unfolding, is a measure of the potency of the denaturant, here FA [46]. For treatment with proteinase K (ProtK), the protein mass in each pellet was first determined by treating one protein pellet with 100% FA and running this sample on SDS-PAGE together with samples of known concentrations of 0.1, 0.2, 0.5 and 0.8 mg/mL. Pellets were then treated with different concentrations of ProtK in a buffer containing 60 mM Tris and 5 mM CaCl2 (pH 8) and incubated at 37°C for 30 min before the reactions were stopped by adding concentrated FA (giving a final [FA] of 50%, equal to 13.25 M). Samples were lyophilized in a Scanvac Coolsafe freeze dryer (Labogene) and subsequently analyzed on SDS-PAGE.

### Cross-interactions between microbiome proteins and αSN

1 mg/mL αSN was fibrillated in the presence of Pr12 or Pr17 seeds using 40 µM ThT in a 96-well plate. A 3 mm glass bead was added to each well, and the program used allowed 300 rpm orbital shaking with readings every 1070 s and 600 s shaking between readings. Pr12 and Pr17 seeds were prepared by incubation overnight at 37°C with slow shaking (300 rpm). Aggregates were then spun down (13,500 rpm, 15 min) and washed twice with MQ before aliquoted into new tubes. Aggregates were pelleted again, and all supernatant was removed. Concentration in the pellet was determined in the same was as described above. To prepare seeds, aggregates of Pr12 and Pr17 were sonicated for 3*10 s with 10 s on ice in between using a QSonica Sonicator and subsequently added at 10%, 5% or 1% (mass/mass) to 1 mg/mL monomeric αSN.

### Transmission Electron Microscopy (TEM) analysis of protein fibrils

The same Pr12 and Pr17 aggregates used to study aggregate stability were also imaged with TEM. Pellets were resuspended in MQ and images were recorded as described [47].

### Fourier transform infrared (FTIR) spectroscopy of protein fibrils

FTIR spectra were recorded on a Tensor 27 instrument (Bruker Optics). 2 µl sample was deposited on the surface of an attenuated total reflection crystal and dried with nitrogen gas. The system was continuously purged with nitrogen gas. Background and water vapor subtractions were performed to obtain a straight baseline. The samples were analyzed with OPUS 5.5. Only the amide I region (1600- 1700 cm^-1^) was used for analysis.

## RESULTS

### Identification of potential amyloid protein candidates in the rat microbiome

Several studies have suggested that the human microbiome is changed in PD patients compared to healthy controls [48–58]. The object of this study was not to attempt to elucidate statistically significant differences between healthy and PD-afflicted rats, but rather to carry out a more general assessment of the occurrence of functional amyloid in the microbiome. Nevertheless, given the potential involvement of the microbiome in the development of PD, we decided to compare healthy rats with a rat model of PD that overexpresses full-length human αSN and develops synucleinopathies both in the CNS and the PNS [32]. Microbial community profiling using amplicon primers targeting the V1-V3 variable region of the 16S rRNA gene revealed an increased abundance of Lachnospiraceae in the PD rats, whereas Lactobacillaceae dominated the WT rats (**Fig. 1A**). However, the relative abundance of most other taxa varied greatly between replicates. Principal coordinate and ANOSIM analyses were carried out on the exact amplicon sequence variants (ASVs) to investigate if there were a statistical difference between the WT and PD rats (**Fig. S1**). A separation along the first principal coordinate (PCo1) was observed between the WT and PD samples. However, the separation was not significant according to the permutations tests on the ordinated data (R=0.7037; *p*-value=0.1; 719 permutations), reflecting the small sample size. Nevertheless, the samples provided an opportunity to search for possible FuBA through the differential solubilization of proteins at higher concentrations of formic acid, as described in Materials and Methods.

**Figure 1.**
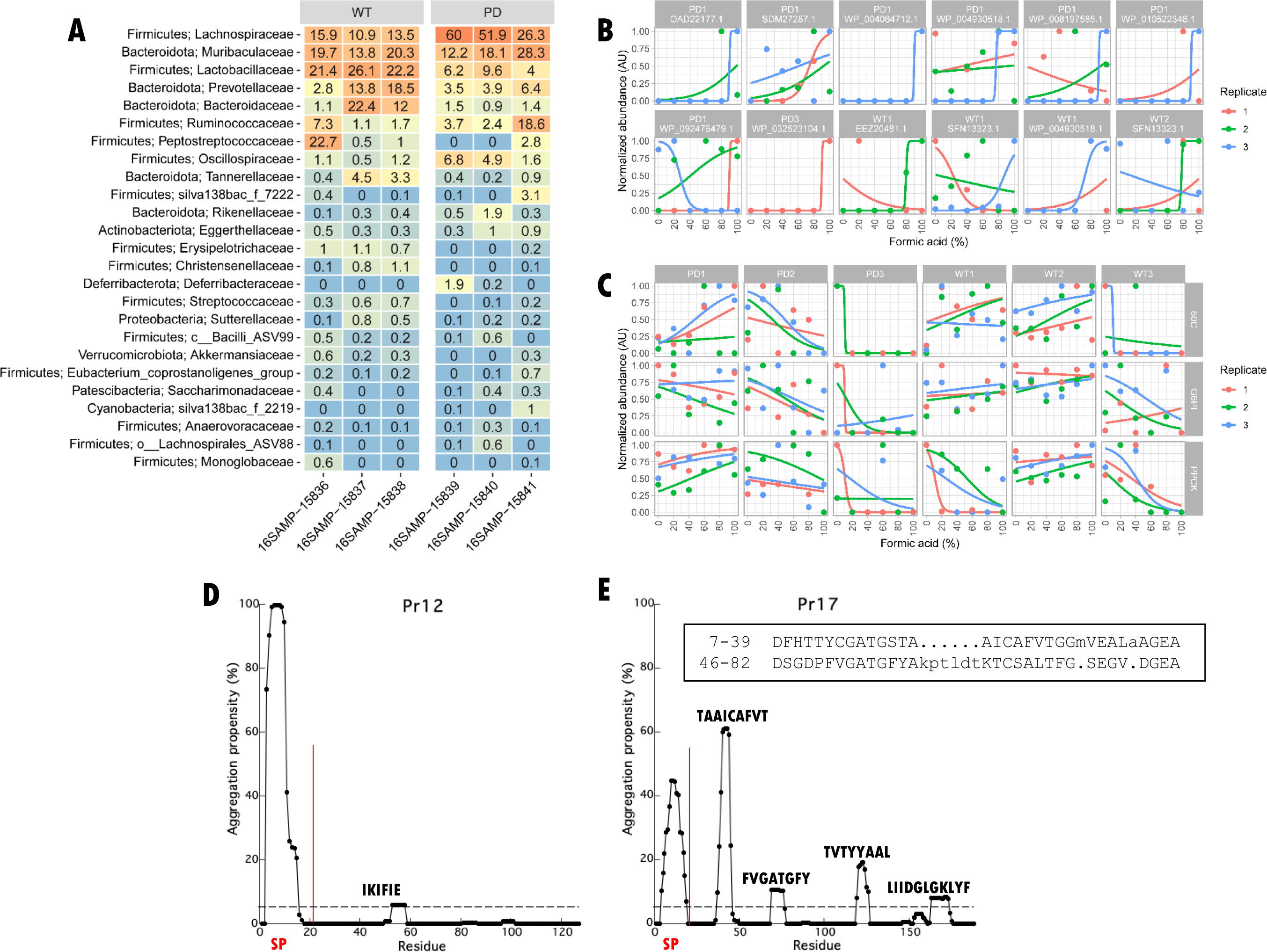
Identification of potential amyloid protein candidates. (**A**) Percent relative abundance of the top 25 bacterial families in wild type (WT) and Parkinson’s disease (PD) rats. The data is based on V1-V3 ASVs classified based on the AutoTax processed SILVA 138 SSURef Nr99 database [36]. Phylum names are provided together with the family names. (**B**) The eleven hit proteins proposed to contain a secretion signal peptide (there are 12 panels because SFN13323.1 is found in both WP1 and WT2). Data points can be hidden behind each other in some of the graphs. Not all proteins were identified in all three replicates. (**C**) Examples of negative controls including the household proteins glycose-6-phosphate isomerase (G6PI), 60 kDa chaperonin (60C) and phosphoenolpyruvate carboxykinase (PPCK). (**D**) TANGO analysis of Pr12 and (**E**) Pr17. The vertical red lines show the location of the signal peptide (SP). The horizontal dashed line represents an aggregation propensity value of 5%. In addition, both protein sequences were analysed with the RADAR tool [43] but imperfect repeats could only be identified for the Pr17 sequence (insert in (E)).

Protein identification with MaxQuant resulted in the identification of between 447 (sample WT3) and 1314 (sample PD2) proteins in each of the six samples (**Table S1**). The proteins were quantified using the LQF algorithm, and their stability towards formic acid was determined as described [31]. Comparing all identified proteins from the WT samples with those identified in the PD samples showed no obvious distinction between the two phenotypes. Moreover, the LQF results were characterized by relatively low reproducibility, as a large fraction of proteins were not identified in all replicates, probably due to low abundance (**Fig. S2**). As the protocol for identification of putative functional amyloids requires a large number of replicates with a sigmoidal signature [31], comparison between WT and PD samples was inconclusive. Therefore, we decided to manually inspect the data and hand-pick the most probable amyloid candidates based on their stability towards formic acid and ignore the number of replicates by which they were identified. We identified between 22 (sample PD3) and 110 (sample PD1) amyloid candidates based on their characteristic sigmoidal signature in plots of normalized protein abundance versus FA concentration (**Fig. 1B** and **Table S1**). Of these 365 proteins, 27 had at least two replicates showing a sigmoidal signature (**Table S2**). One particularly interesting protein is elongation factor Tu (EF-Tu) which appears three times in the PD1 sample. In fact, EF-Tu appears as a hit 46 times (although in most of the cases only a single replicate shows the sigmoidal signature) across all samples and equally divided between PD (22 times) and WT samples (24 times) (data not shown). EF-Tu is interesting because the *Gallibacterium anatis* EF-Tu (ID: KGQ60852.1) was recently identified as a functional amyloid-forming protein [59], indicating that this FA-approach is able to identify amyloid-relevant proteins. Household proteins glucose-6-phosphate isomerase (G6PI), 60 kDa chaperonin (60C) and phosphoenolpyruvate carboxykinase (PPCK) – which were identified 71, 38 and 143 times, respectively, across all 6 samples – generally do not show any specific signature across the FA concentration range (**Fig. 1C**) (though a few peptides of PPCK were identified to follow a sigmoidal behavior and consequently led to the protein’s inclusion in Table S1).

Because many functional amyloid systems are dependent on secretion through the Sec translocon, all protein hits were analyzed with the three different signal peptide prediction tools SignalP 4.1 [42], DeepSig [60] and SignalP 5 [61] and ranked according to the probability of containing a signal peptide. From this analysis 11 proteins were identified (**Table 1**). Out of these 11 proteins, only three proteins were predicted to contain a signal peptide with all three tools. We decided to select two proteins for recombinant expression to investigate their amyloidogenicity experimentally. We chose to limit ourselves to proteins predicted to contain a signal peptide by all 3 signal peptide predictors and to exclude large (> ca. 450 residues) proteins to avoid low recombinant expression levels. This left proteins WP_032523104.1 (12 kDa without the 22 aa signal peptide) and OAD22177.1 (17 kDa without the 18 aa signal peptide) which we will from now on refer to as Pr12 and Pr17, respectively. Pr12 was identified in sample PD3 while Pr17 was found in sample PD1. According to the Uniprot database Pr12 is a DUF1499 domain-containing protein produced by the cyanobacterium *Prochlorococcus marinus* while Pr17 is a secreted protein produced by the vacuolate sulfur bacteria *Candidatus Thiomargarita nelsonii*. To obtain residue-specific information on the potential aggregation of Pr12 and Pr17, we used the TANGO web server software [62] to predict aggregation-prone regions. For both proteins it was seen that the signal peptide (SP) gave a high aggregation propensity (**Fig. 1DE**), but interestingly, Pr17 also showed four separated stretches of 8- 12 residues with high aggregation propensity (aggregation propensity value > 5%, dashed line) throughout the proteins sequence (**Fig. 3b**). Both proteins were also investigated for the presence of imperfect repeats using the RADAR tool [43]. Whereas no repeats could be found for Pr12, two repeats were identified in the Pr17 sequence (Fig. 3b, insert). The repeats showed low conservation but, interestingly, two of the highly amyloidogenic stretches (TAAICAFVT and FVGATGFY) were covered by the repeat sequences.

**Table 1:** 11 proteins from the PD microbiome containing at least two replicates with sigmoidal solubility curves and predicted to have a signal peptide.

| Protein ID | Software with a positive signal peptide prediction | Mature protein length (aa) |
| --- | --- | --- |
| OAD22177.1 (Pr17) | DeepSig, SignalP 4.1, SignalP 5 | 170 |
| SDM27287.1 | DeepSig, SignalP 4.1, SignalP 5 | 480 |
| WP_032523104.1 (Pr12) | DeepSig, SignalP 4.1, SignalP 5 | 105 |
| WP_004064712.1 | SignalP 4.1, SignalP 5 | 1475 |
| WP_010522346.1 | SignalP 4.1, SignalP 5 | 808 |
| EEZ20481.1 | SignalP 5 | 380 |
| SFN13323.1 | SignalP 4.1 | 411 |
| SFR13334.1 | SignalP 5 | 361 |
| WP_004930518.1 | SignalP 4.1 | 172 |
| WP_008197585.1 | SignalP 5 | 121 |
| WP_092476479.1 | SignalP 4.1 | 537 |

### Pr12 and Pr17 both form ThT-positive aggregates with amyloid characteristics

It was possible to recombinantly produce both proteins in *E. coli* at yields of 3.5-4.5 mg/L culture. The proteins were then purified to satisfying purity, as visualized by SDS-PAGE, by Ni-NTA chromatography via a C-terminal His6 tail (**Fig. S3**). To analyse the two protein’s propensity to form amyloid, we started by analysing their behaviour in a standard ThT fibrillation assay. ThT is a dye commonly used to study amyloid fibrillation as it shows a bright fluorescence when bound the amyloid fibrils and therefore is a suitable reporter for fibril formation. After desalting into PBS, the concentration of the two proteins was measured and aggregation of the two proteins was immediately investigated with ThT. Note that when measuring protein concentration directly after desalting, the protein solution showed light scattering (tailing of the protein peak at wavelengths > 320 nm), indicating protein aggregation (data not shown). An instant increase in ThT signal could be observed for both proteins (**Fig. 2AB, left panels**) but where Pr17 reaches a stable end level, the signal for Pr12 starts decreasing after reaching a maximum intensity after ∼ 6 hours. No lag phase (which is otherwise usually seen during protein fibrillation) was observed for either protein. We note a short dip in the fluorescence signal around t = 0 (**Fig. S4**) which we ascribe to fluorescence quenching as the temperature increases to 37°C. FTIR analysis of the resulting fibrils showed a pronounced peak around 1622 cm^-1^ for Pr12 and between 1624-1627 cm^-1^ for Pr17 (**Fig. 2 right panel**). Amyloid fibrils normally absorb strongly in the 1615-1630 cm^-1^ region [63] while native β-sheets absorb at wavenumbers in the 1630-1643 cm^-1^ range [64]. This indicates that Pr12 and Pr17 aggregates are amyloid-like.

**Figure 2.**
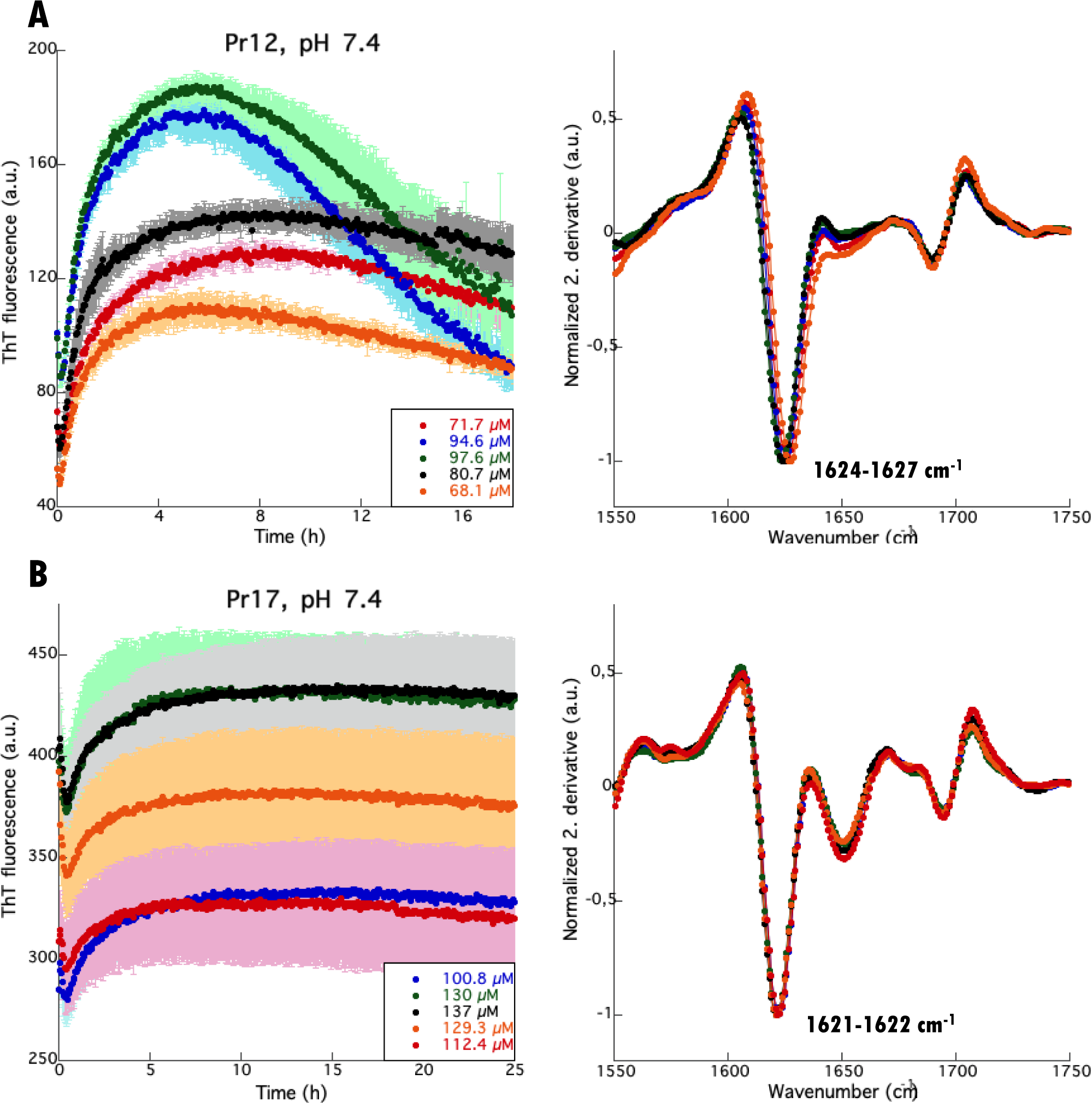
Pr12 and Pr17 readily form ThT-positive aggregates. (**a**) Pr12 and (**b**) Pr17 were immediately subjected to a ThT fibrillation assay (left panel) after desalting. FTIR (right panel) was performed on the pellet after fibrillation and showed cross-β characteristic peaks between 1620-1630 cm^-1^.

### Fibrillation of Pr12 is sensitive to urea

To avoid premature aggregation of the proteins, the denaturant urea was added to the desalting buffer. The two proteins differed somewhat in their urea sensitivity. While ≥ 1.5 M urea completely inhibited aggregation of Pr12 (**Fig. 3A**), 0.5 mg/ml Pr17 maintained aggregation up to 2.5 M urea (**Fig. 3C**). Following fibrillation, the samples were spun down and the supernatants were analysed with SDS-PAGE. For Pr12, a good correlation was seen between the samples showing aggregation (0.2 M, 0.5 M, and 1 M urea) and the loss of protein from the supernatant (**Fig. 3B** and **Fig. S5**). However, for Pr17 all protein stayed in solution after centrifugation (data not shown) despite the increase in ThT fluorescence seen for the 0.4-2.5 M urea samples. To find out whether the difference in ThT fluorescence was caused by the formation of different species of Pr17, we analysed these samples with SEC. The results confirmed that this was the case: Pr17 samples giving high ThT fluorescence signals (0.4-2.5 M urea) contained higher order oligomers that eluted from the column together with the void volume, while samples with low/no ThT fluorescence signal (4 and 6 M urea) only show the Pr17 monomer peak eluting around 17 mL (**Fig. 3D**). We ascribe the shift in monomer elution peak position with rising [urea] to either increased expansion of the protein or reduced interactions with the column. This earlier elution is even more pronounced for free ThT which elutes around 37 ml in 0.4M urea but around 24 ml in 6M urea (**Fig. S6A**). Interestingly, ThT elution volume and end point ThT levels of Pr17 decline in a similar fashion as a function of [urea] (**Fig. 3E**, red curve). Plotting the monomer/oligomer intensity as a function of urea concentration shows a cross-over (*i.e.* roughly equal amounts of both species) at 2.25 M urea (**Fig. 3F**).

**Figure 3.**
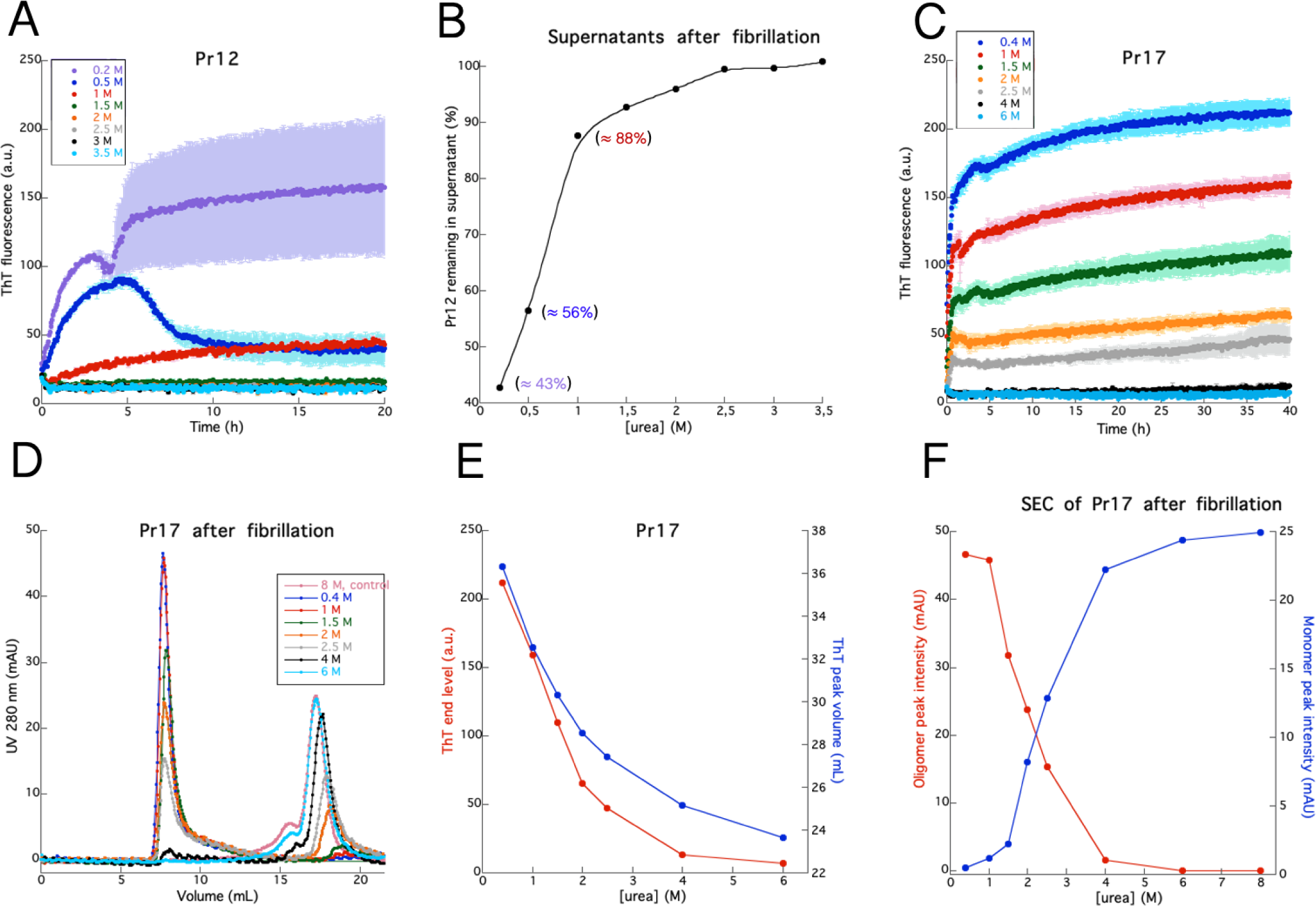
Fibrillation of Pr12 and Pr17 in the presence of urea. (**A**) Investigation of Pr12 aggregation in the presence of increasing concentrations of urea. (**B**) Amount of soluble protein left after aggregation determined by SDS-PAGE of the supernatants after centrifugation of samples in (A). (**C**) Investigation of Pr17 aggregation in the presence of increasing concentrations of urea followed by (D) SEC analysis of the same samples including a control of Pr17 in 8 M urea (to keep it monomeric). (E) Correlation between the ThT end levels from (C) and the ThT elution volumes from (D). (**F**) Correlation between the decrease in Pr17 oligomer and the increase in monomeric Pr17 with increasing urea concentrations.

Pr17 contains four Cys residues (position 32, 42, 87 and 103). SDS-PAGE of the SEC samples under both non-reducing (**Fig. S6B**) and reducing (**Fig. S6C**) conditions revealed a disulphide-bonded dimer band under non-reducing conditions (absent under reducing conditions), which is also seen in SEC as a shoulder to the monomer peak (**Fig. 3D**). Oligomers form a smear in the top part of the non-reducing gel. Note that the amount of monomer is essentially constant over 0.4-8M urea on the reducing gel. This indicates that oligomers and dimers dissociate to monomers when exposed to SDS and reducing conditions.

### pH dependence of Pr12 and Pr17 fibril formation

We also investigated the pH sensitivity of the two proteins’ aggregation in view of the variable pH environment encountered in the GI tract. Pr12 showed a rapid increase in fluorescence within a few hours, particularly between pH 4-7.4, but again no lag phase could be observed (**Fig. 4A**). Fibrillation was essentially abolished at pH 3 and pH 8. At pH 7.4, the fibrillation curve showed an instant increase followed by a decrease after 5 hours. We tested if this was an effect of the buffer by carrying out fibrillation in 0-3.5M urea with 60 mM Tris buffer instead of PBS at pH 7.4 but still observed the same ‘overshoot’ behaviour (**Fig. S7A**) though with significant variation between the triplicates (**Fig. S7B**). The solubilisation of Pr12 at higher [urea] in Tris buffer (**Fig. S7C**) was essentially identical to that in PBS buffer (**Fig. 3B**). A 7-fold replication of aggregation of 0.5 mg/mL Pr12 in 60 mM Tris, pH 7.4 showed variation largely in their ThT endpoint (**Fig. S7D**) although FTIR analysis showed identical secondary structure (**Fig. S7E**). We therefore conclude that the ‘overshoot’ behaviour reflects a stochastic aggregation behaviour at this pH to form different levels of higher-order aggregates (with consequent different levels of precipitation and light-scattering). This indicates that Pr12 has a different aggregation mechanism at neutral pH compared to both more basic and more acidic conditions where the aggregation curves are more reproducible. Expasy predicts Pr12 to have a theoretical pI of 4.98 which cannot account for the big difference seen between pH 6 and pH 7.4.

**Figure 4.**
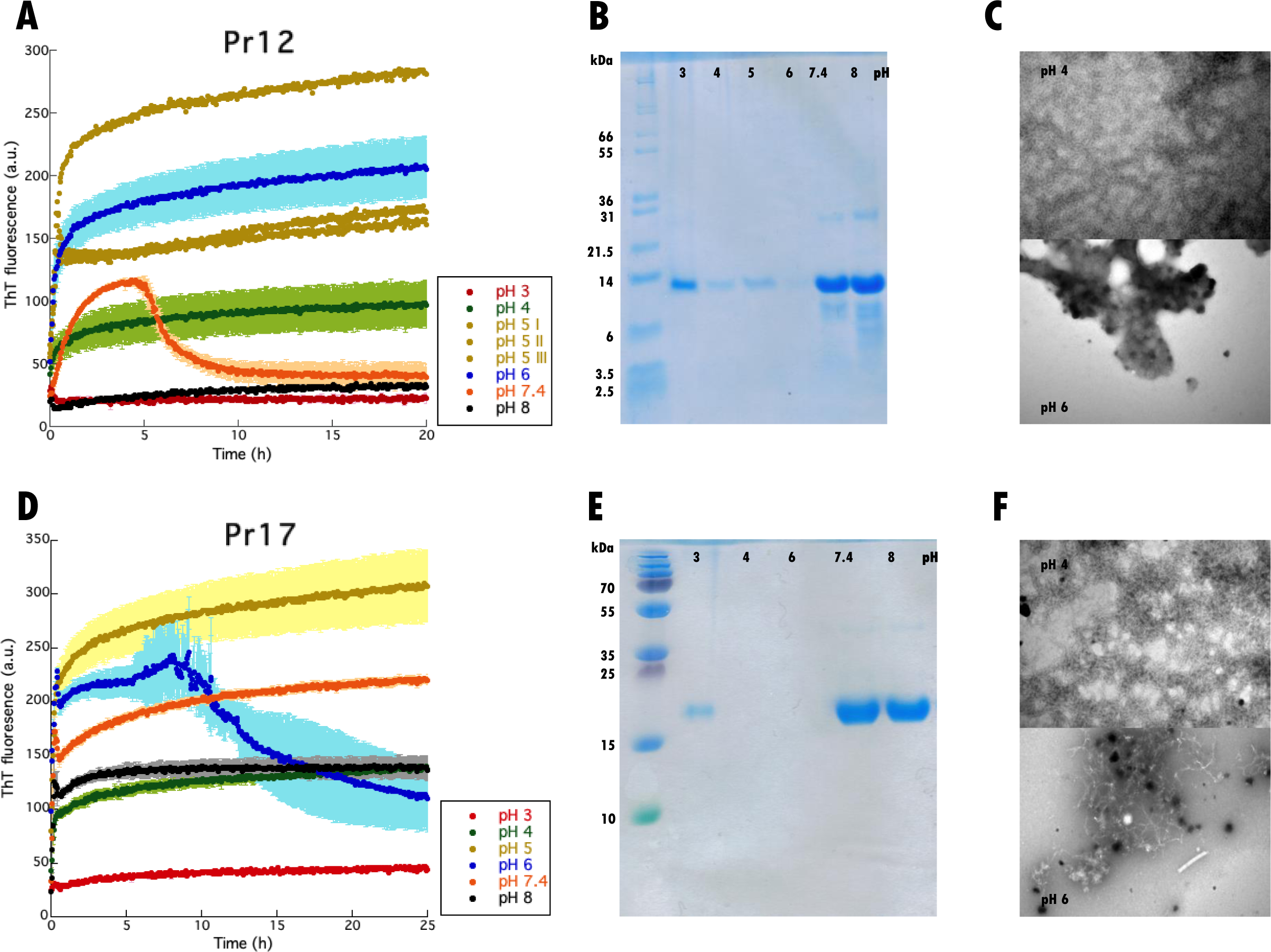
pH dependence of Pr12 and Pr17 aggregation. Pr12 (**A**) and Pr17 (**D**) were aggregated in the presence of ThT at different pH values and the samples were afterwards separated into soluble and insoluble fractions by centrifugation. The soluble part (the supernatant) was analysed with SDS- PAGE for Pr12 (**B**) and Pr17 (**E**). TEM images of Pr12 (**C**) and Pr17 (**F**) aggregates formed at either pH 4 (top) or pH 6 (bottom).

Pr17 aggregation is relatively robust to pH changes and happens at all pH values tested, although only to a very small extent at pH 3 (**Fig. 4D**). The theoretical pI of Pr17 is 4.27. At pH 6, the aggregation curves always showed great variation between different experiments and sometimes gave an overshoot (**Fig. 4D**) but with great variation between individual runs (**Fig. S8**).

The extent of aggregation was measured by SDS-PAGE of soluble (supernatant) and insoluble (pellet) protein after the ThT assay both for Pr12 (**Fig. 4B**) and Pr17 (**Fig. 4E**). Pr12 showed very low levels of soluble protein between pH 4 and 6, corresponding nicely with the high ThT end levels. At pH 7.4 and 8 the protein stays soluble despite the initial increase in ThT seen at pH 7.4. Similarly, for Pr17 after 25 hours fibrillation, we saw very low solubility at pH 3, 4 and 6 and much higher solubility at pH 7.4 and pH 8 despite the fact that the end-point ThT levels are identical for pH 4 and pH 8. TEM analysis (**Fig. 4C** and **4F**) also showed pH-dependent differences in aggregate structure: at pH 4, both proteins formed a mesh of very thin, fibrillar structures while at pH 6, Pr12 showed more amorphous aggregates with no apparent fibrils being present and Pr17 showed a mix of fibrils and amorphous structures.

### Pr17 aggregates are more stable than Pr12 towards urea and FA but not proteinase K

The high stability of the FuBA structure ensures that functional amyloids stay insoluble under denaturing conditions and only dissociate into monomers at high concentrations (80-100%) of FA [29, 30, 65]. We therefore decided to investigate whether Pr12 and Pr17 aggregates show the same degree of stability towards denaturants like urea and FA. Aggregates were mixed with increasing concentrations of either urea or FA and the supernatant analyzed with SDS-PAGE (**Fig. 5AB**). By normalizing to the 100% FA band (where we assume 100% of the fibrils dissolve) we can determine the denaturant concentration required to dissolve half of the fibrils, [den]^50%^. For Pr17 we determined [urea]^50%^ to be ∼ 4.4 M (as determined by visual inspection) and [FA]^50%^ to be 32.7 ± 3.7% after fitting the data points to a sigmoidal curve. Pr12, in contrast, was highly sensitive towards FA with a [FA]^50%^ around 0.1% (**Fig. 5B**). The [FA]^50%^ value of ∼ 30% for Pr17 is only slightly lower than the stabilities of [FA]^50%^ of around 50% we normally see for functional amyloids [66]. In contrast, the [FA]^50%^ value of 0.1% for Pr12 is similar the [FA]^50%^ value determined for αSN fibrils [66]. Thus *in vitro* aggregates of Pr17 show aspects of a classical functional amyloid protein while Pr12 aggregates appear much less stable.

**Figure 5.**
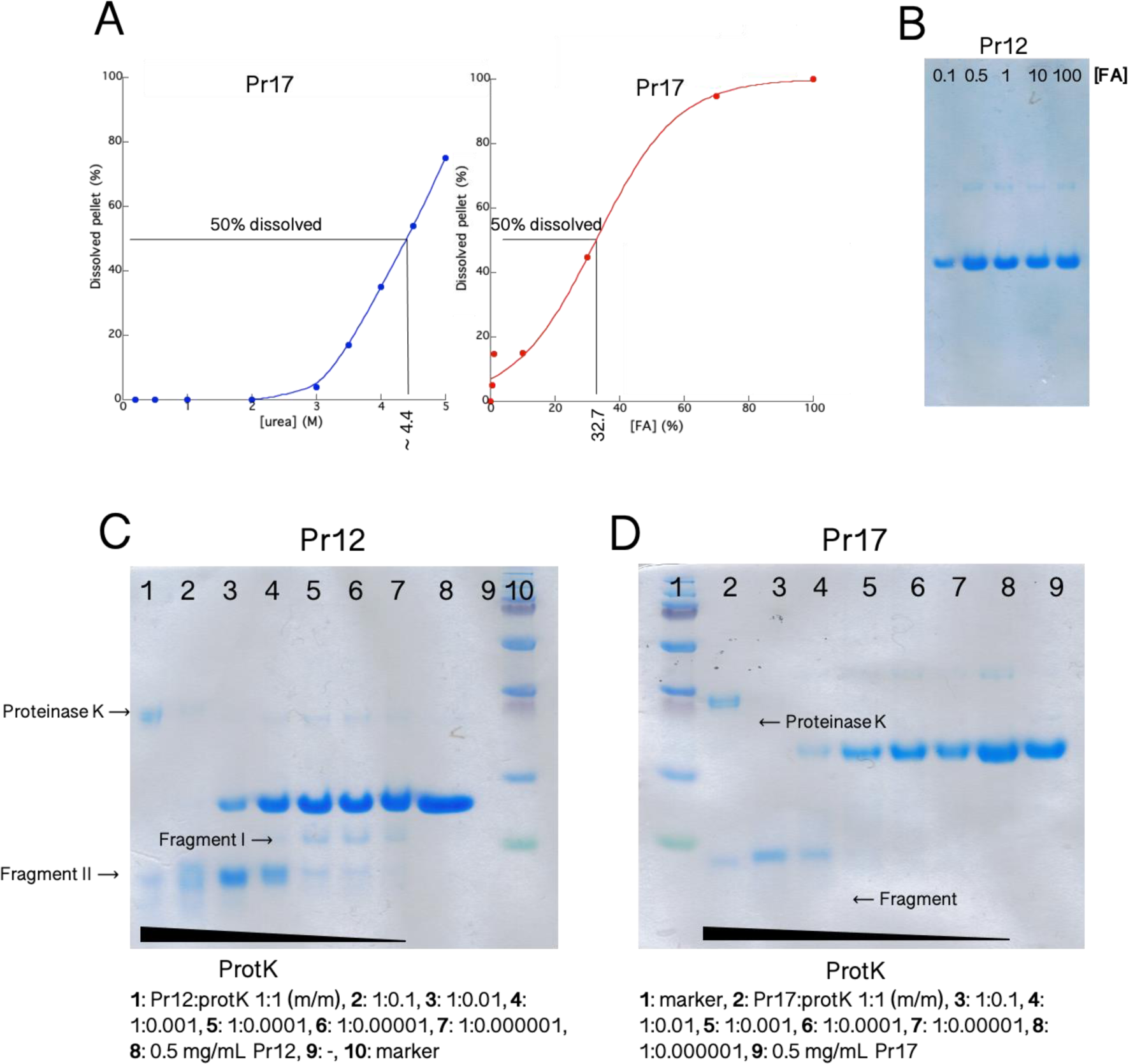
Stability investigations of Pr12 and Pr17 aggregates. (**A**) Aggregates formed by Pr17 were dissolved in increasing concentrations of urea (left) or formic acid (right) and the amount of dissolved protein was visualized using SDS-PAGE after lyophilization. Quantification was performed on these gels using ImageJ. (**B**) Aggregates formed by Pr12 were dissolved in increasing concentrations of formic acid and dissolved protein was visualized with SDS-PAGE after lyophilization of 20 µL of the supernatant. The Pr12 aggregates dissolved at very low formic acid concentrations. Aggregates of Pr12 (**C**) and Pr17 (**D**) were subjected to digestion with increasing ratios of protein:proteinase K up to 1:1 (m/m) followed by SDS-PAGE analysis.

We also treated protein aggregates with increasing concentrations of proteinase K (ProtK) to elucidate if the aggregates formed by the proteins – especially the more stable Pr17 aggregates – contained a protected core. In this case, we would expect to see a clear band on SDS-PAGE after ProtK treatment. High concentrations of proteinase K (Pr12/Pr17:protK ratios of 1:1 (m/m) and 1:0.1 (m/m)) resulted in a break-down to smaller fragment of ∼ 7 kDa for Pr12 (**Fig. 5C**, fragment II) and ∼ 9 kDa for Pr17 (**Fig. 5D**) at sample/ProtK mass ratios down to 1:0.01 (lane 4). This indicates a structured core present in both protein aggregates. At sample:protK (m/m) ratios of 1:0.001, appr. 40% of Pr12 (Fig. 9c, lane 3) and 15% of Pr17 (Fig. 9d, lane 4) remained full-length after treatment. Interestingly, for Pr12 a slightly bigger fragment of ∼ 11 kDa appeared when treated with lower proteinase K concentrations (**Fig. 5C**, lane 4-7). However, the full-length protein still dominated. This 11 kDa fragment is the first to be cleaved off (but only to a low degree as no apparent change can be seen in the intensity of the full-length protein band) by ProtK and is fully degraded at higher ProK concentrations. The 9 kDa fragment of Pr17 appears to be relatively protected from the proteolytic attack when in the aggregated structure. However, the intensity of the band is lower at the 1:1 ratio (lane 2) than the 1:0.1 ratio (lane 3), so it is not completely resistant towards proteolytic degradation.

### Co-incubation of Pr12 and Pr17 with αSN

To study the effects of Pr12 and Pr17 on fibrillation of αSN, we compared the fibrillation of monomeric αSN in the absence and presence of these two proteins. We incubated monomeric αSN with seeds made from aggregated Pr12 or Pr17. As seen in **Fig. S10**, neither Pr17 seeds (left) or Pr12 seeds (right) had any significant effect on αSN fibrillation which in all cases fibrillated with a lag phase of ∼ 5 hours.

## DISCUSSION

### Bacterial biofilm in the GI as a source of functional amyloid

Functional amyloids such as curli fibrils are often part of bacterial biofilms. Up to 40% of all bacteria form biofilms [67] and it therefore comes as no surprise that biofilms and bacterial microcolonies have been observed in the human GI tract [68–70] where up to 10^11^ colony forming units (CFU) are found per mL gut content in the colon [71]. In addition, bacterial biofilms have been found associated to food particles in faecal samples from healthy individuals [27, 72] and have also been visualized directly in mice [73]. To date, however, studies of mucosal biofilms in the GI tract are largely limited to diseases like ulcerative colitis [74, 75], peptic ulcer disease [76], Crohn’s disease and cases of diarrhea [77].

Here we sought to identify amyloid proteins in the microbiome of PD Tg rats and their healthy WT counterparts. One aspiration was to establish if the PD microbiome was enriched for amyloidogenic proteins, which might in turn indicate that microbial products produced in the GI tract could play a role in neurodegenerative diseases like Alzheimer’s [78] and Parkinson’s disease [12, 79] as suggested. However, we observed strikingly similar numbers of hit proteins identified from either of the two groups (181 hits for the three PD samples vs 186 hits for the WT) and therefore cannot make any definite conclusions in this regard. Two proteins, here referred to as Pr12 and Pr17, were identified based on a combination of experimental (formic acid solubility/MS), bioinformatics analysis (the presence of a signal peptide and high amyloidogenicity across the amino acid sequence) and feasibility (a modest size compatible with recombinant expression). Both proteins were originally identified from the PD samples (Pr12 from the PD3 sample and Pr17 from the PD1 sample). The two proteins have not been biophysically characterized before but we show that both proteins readily form ThT-positive aggregates to some extent at all pH values investigated (pH 3-8). Pr17 even aggregates in the presence of high concentrations of urea. The aggregation process, however, did not follow classical sigmoidal kinetics which involve an initial lag phase where nuclei are formed and grow [80]. The absence of a lag phase is indicative of amorphous aggregation, even though fast amyloid formation has also been reported [81, 82]. The aggregates formed at pH 7.4 for both proteins show a major band between 1621 and 1627 cm^-1^ when analysed with FTIR, indicating extended β-sheet structures like those present in amyloid fibrils. TEM images also revealed worm-like fibril structures for both proteins. However, when the stability of the aggregates towards FA was investigated, Pr12 showed extreme sensitivity and dissolved at very low FA concentrations ([FA]^50%^ ∼ 0.1%) while Pr17 aggregates were more robust and required 32.7% FA for half of the aggregated material to dissolve.

This value is lower than the 80-100% required to dissolve functional amyloid fibrils such as CsgA and FapC into protein monomers [29, 65] (and also lower than the 50% value used as a criterion when searching the MS data for possible hits, cf. Materials and Methods and [31]). However, other functional amyloids, such as the TasA fibrils produced by *Bacillus subtilis*, have also been found to dissolve at lower (∼ 20%) FA concentrations [83] and we therefore cannot exclude that a given protein is an amyloid based on the FA stability alone. The presence of several amyloidogenic hotspots and two imperfect repeats in the Pr17 sequence further support its robust fibrillation behaviour.

### pH sensitivity of fibrillation and the GI tract

It is noteworthy that Pr12 and Pr17 behave differently in terms of forming soluble vs insoluble aggregates between pH 3 and 8 in view of the great variation of intraluminal pH throughout the GI tract [84]. In the stomach, the pH can be as low as 1.5 in the fasting state (though food consumption increases this within minutes to around pH 5) [85] but rapidly changes to pH 6 in the duodenum and increases further to around neutral at the end of the small intestine. This is followed by a second drop to pH 5.7 in the cecum and a final increase to 6.7 when reaching the rectum [84]. Since the bacterial count in the stomach is very low (< 10^3^ CFU per mL [86]), we believe that the conditions in the small and large intestines (pH values ranging from 5.7 to 7.4) are particularly relevant.

### Perspectives on the role of functional amyloid in PD

Based on the results presented here, we conclude that Pr17 holds great promise as a novel functional amyloid protein while Pr12 instead appears to form less structured β-aggregates. The presence of a relatively protease-resistant aggregation core indicates however that both proteins are able to form regular intermolecular contacts. We speculate that Pr12 might form even more stable amyloid fibrils in the presence of co-factors that could be available *in vivo*. Neither of the two proteins had any seeding effect on αSN fibrillation. This, however, does not necessarily mean that the microbiome is not involved in the development of PD. First of all, the rats investigated were relatively young (3 months) and the signs of disease at this age is still very modest, as evaluated by subtle locomotor deficits, reduced ability to discriminate smells and more dot-like appearance of αSN in the striatum nerve end terminals [32]. At this age, no deficits in locomotor activity could be detected and the amount of insoluble (urea-treated) αSN in most brain regions investigated (olfactory bulb, striatum, substantia nigra and hippocampus) was unchanged [32].

## ACKNOWLEDGEMENTS

D.E.O. acknowledges support from Innovation Foundation Denmark (Grant 5188-00003B) through the Joint Programme on Neurodegenerative Diseases (aSynProtec), the Lundbeck Foundation (Grant R276-2018-671) and the Independent Research Council Denmark, Natural Sciences (Grant 8021- 00208B). M.B. acknowledges support from the National Science Centre grant 2019/35/B/NZ2/03997.

## Supplementary Information

### SUPPLEMENTARY TABLES

**Table S1:** Number of proteins and putative amyloids identified by MS/MS.

|  | PD1 | PD2 | PD3 | WT1 | WT2 | WT3 |
| --- | --- | --- | --- | --- | --- | --- |
| Proteins identified | 1214 | 1314 | 1145 | 1048 | 1183 | 447 |
| Putative amyloids | 110 | 49 | 22 | 89 | 73 | 24 |
| Putative amyloids (%) | 9.1 | 3.7 | 1.9 | 8.5 | 6.2 | 5.4 |

**Table S2:** Proteins with amyloid characteristic formic acid resistance in at least two replicates.

| NCBI accession | Description | Sample |
| --- | --- | --- |
| CCY16765.1 | 30S ribosomal protein S10 | PD1 |
| WP_039646517.1 | MULTISPECIES: 50S ribosomal protein L11 | PD1 |
| WP_057773609.1 | elongation factor Tu<sup>a</sup> | PD1 |
| WP_075126584.1 | elongation factor Tu<sup>a</sup> | PD1 |
| WP_003648636.1 | elongation factor Tu<sup>a</sup> | PD1 |
| WP_093135193.1 | molecular chaperone DnaK | PD1 |
| OFX39478.1 | pyruvate:ferredoxin (flavodoxin) oxidoreductase | PD1 |
| CBL09414.1 | hypothetical protein ROI_24490 | PD1 |
| SCH57546.1 | Pyruvate%2C phosphate dikinase<sup>a</sup> | PD1 |
| WP_004071889.1 | flagellin | PD1 |
| WP_011245737.1 | MULTISPECIES: molecular chaperone GroEL | PD1 |
| WP_016228711.1 | formate acetyltransferase | PD1 |
| WP_084256702.1 | L-rhamnose isomerase | WT1 |
| WP_092969895.1 | sn-glycerol-3-phosphate ABC transporter ATP-binding protein UgpC | WT1 |
| GAP43239.1 | DNA-directed RNA polymerase subunit $\alpha$ | WT1 |
| CDD47242.1 | putative uncharacterized protein | WT1 |
| EQB89936.1 | hypothetical protein M918_18135 | WT1 |
| CDF42960.1 | 4-hydroxy-3-methylbut-2-enyl diphosphate reductase/S1 RNA-binding domain protein | WT1 |
| OGH97713.1 | Reverse rubrerythrin-1 | WT1 |
| WP_069198253.1 | lysine-sensitive aspartokinase 3, partial | WT1 |
| SDA77697.1 | formate C-acetyltransferase | WT1 |
| WP_004055611.1 | ABC transporter ATP-binding protein | WT1 |
| WP_008391483.1 | MULTISPECIES: 50S ribosomal protein L27 | WT1 |
| CCX47505.1 | ATP synthase subunit $\alpha$ | WT1 |
| WP_072742756.1 | F<sub>0</sub>F<sub>1</sub> ATP synthase subunit $\beta$ | WT2 |
| WP_074650570.1 | pyruvate, phosphate dikinase<sup>a</sup> | WT2 |
| WP_009757376.1 | phosphoenolpyruvate carboxykinase | WT2 |

Note:
<sup>a</sup> These proteins, though described by the same name, represent different homologs in the NCBI database used as a reference for peptide mapping.

### SUPPLEMENTARY FIGURES

**Figure S1.**
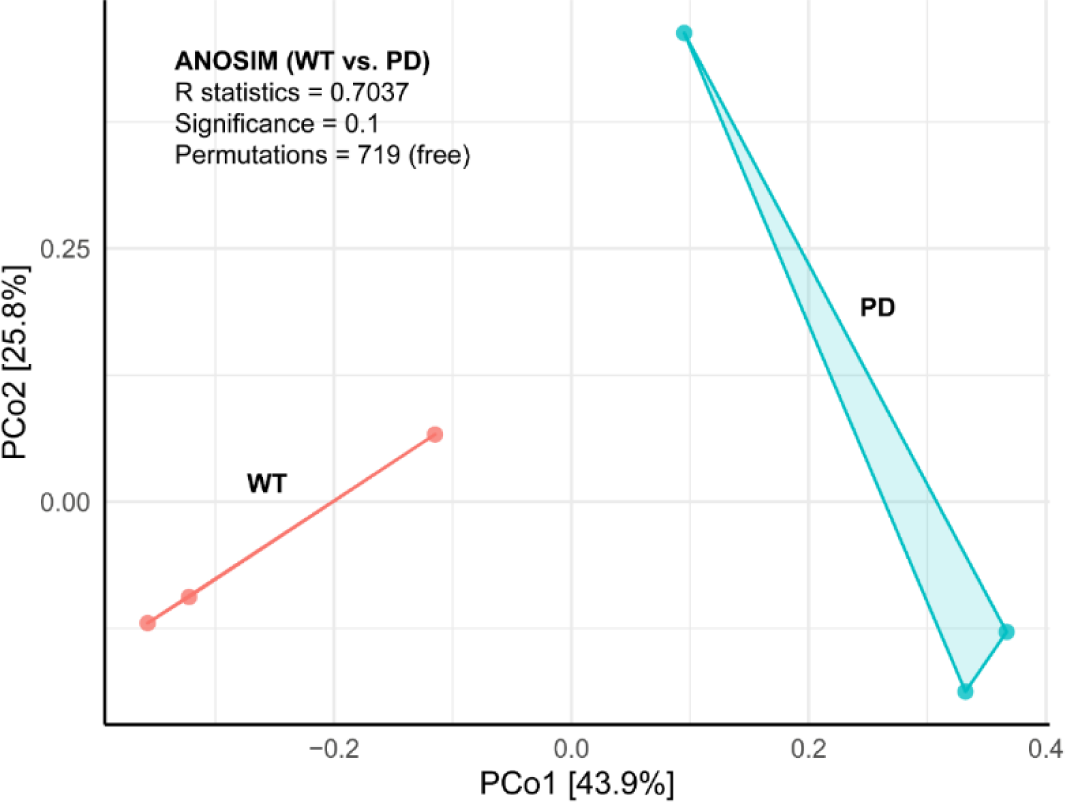
Principal Coordinates Analysis (PCoA) based on the Bray-Curtis distance measure (108) of 6 samples and 1179 ASVs. Prior to the analysis, ASVs that are not present in more than 0.01% relative abundance in any sample were removed. No initial data transformation was applied. The relative contribution (eigenvalue) of each axis to the total inertia in the data is indicated in percent in the axis titles. ANOSIM was used to statistically test whether there was a significant difference in the bacterial composition between WT and PD rats. The statistics are provided in the plot.

**Figure S2.**
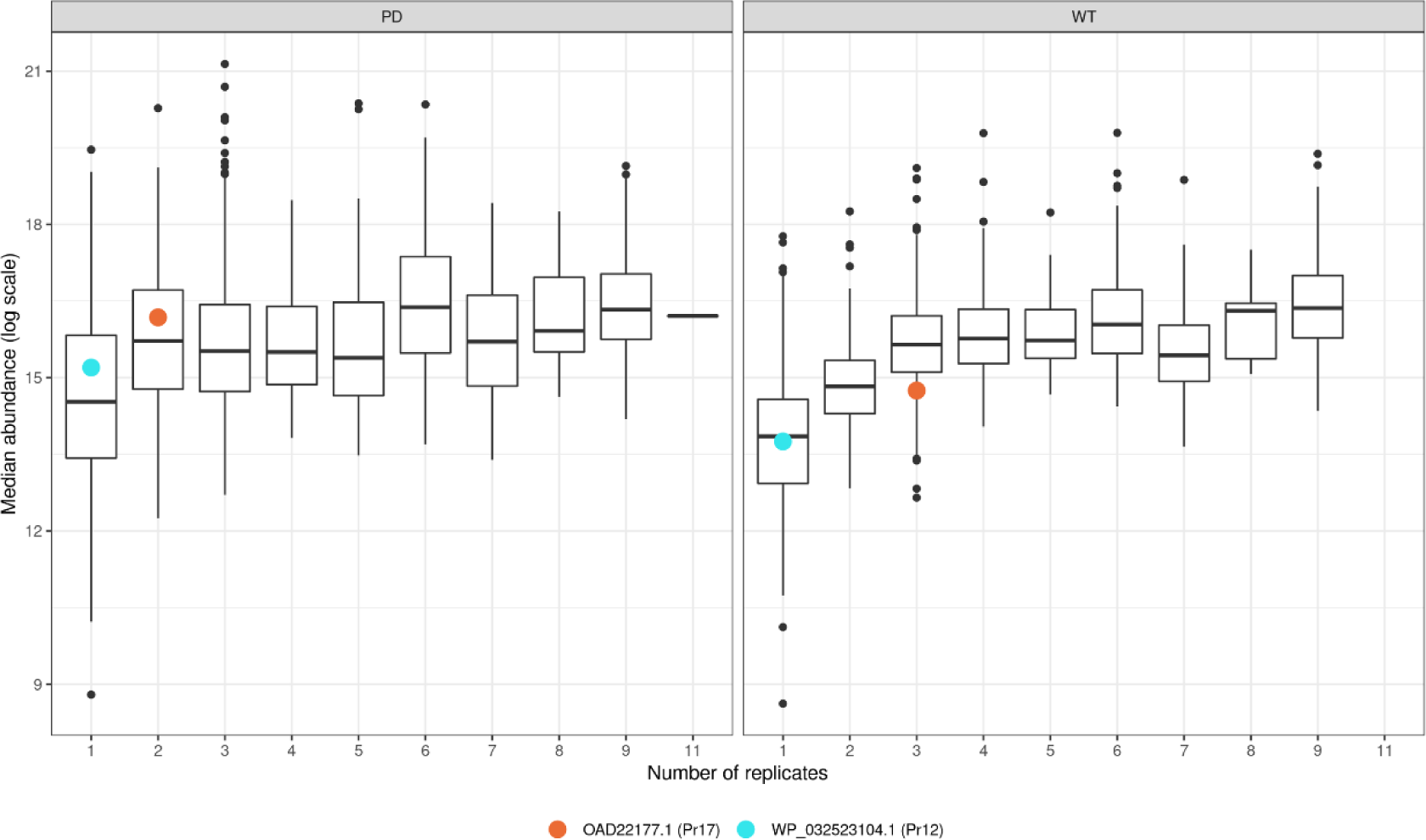
The occurrence of proteins identified by LQF analysis and their median abundance (log-transformed). The x-axis represents the number of replicates where tryptic peptides from a specific protein was identified. The two proteins investigated in detail in this study (Pr17 and Pr12) are shown in orange and blue, respectively.

**Figure S3.**
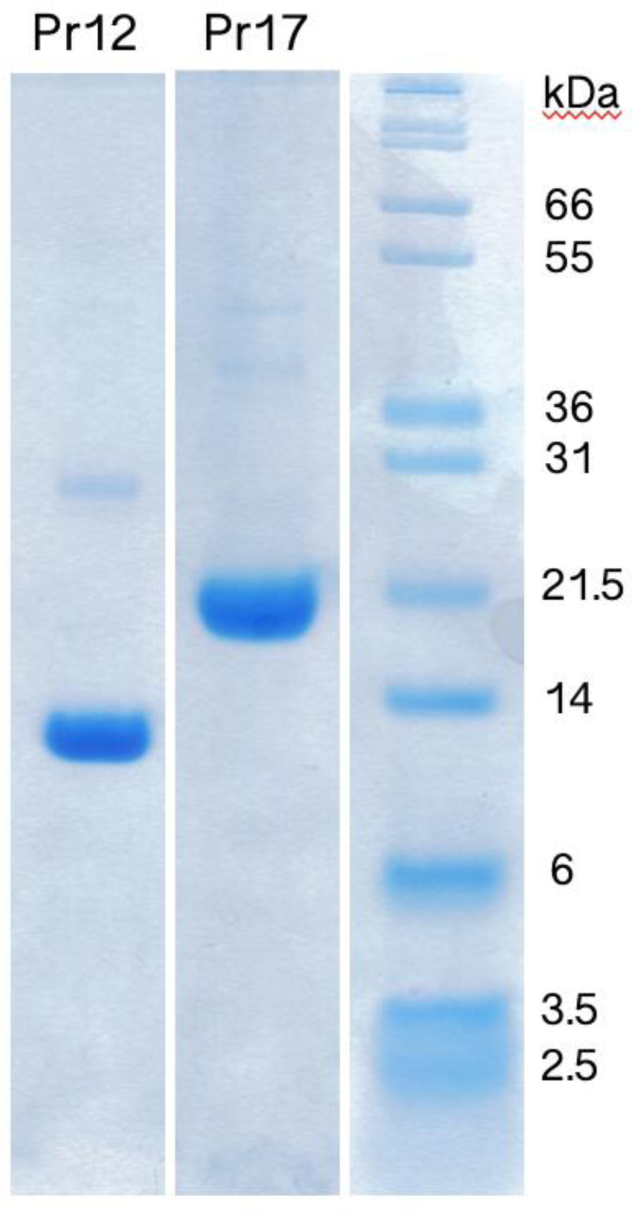
SDS-PAGE analysis of Ni-NTA purified samples of Pr12 and Pr17.

**Figure S4.**
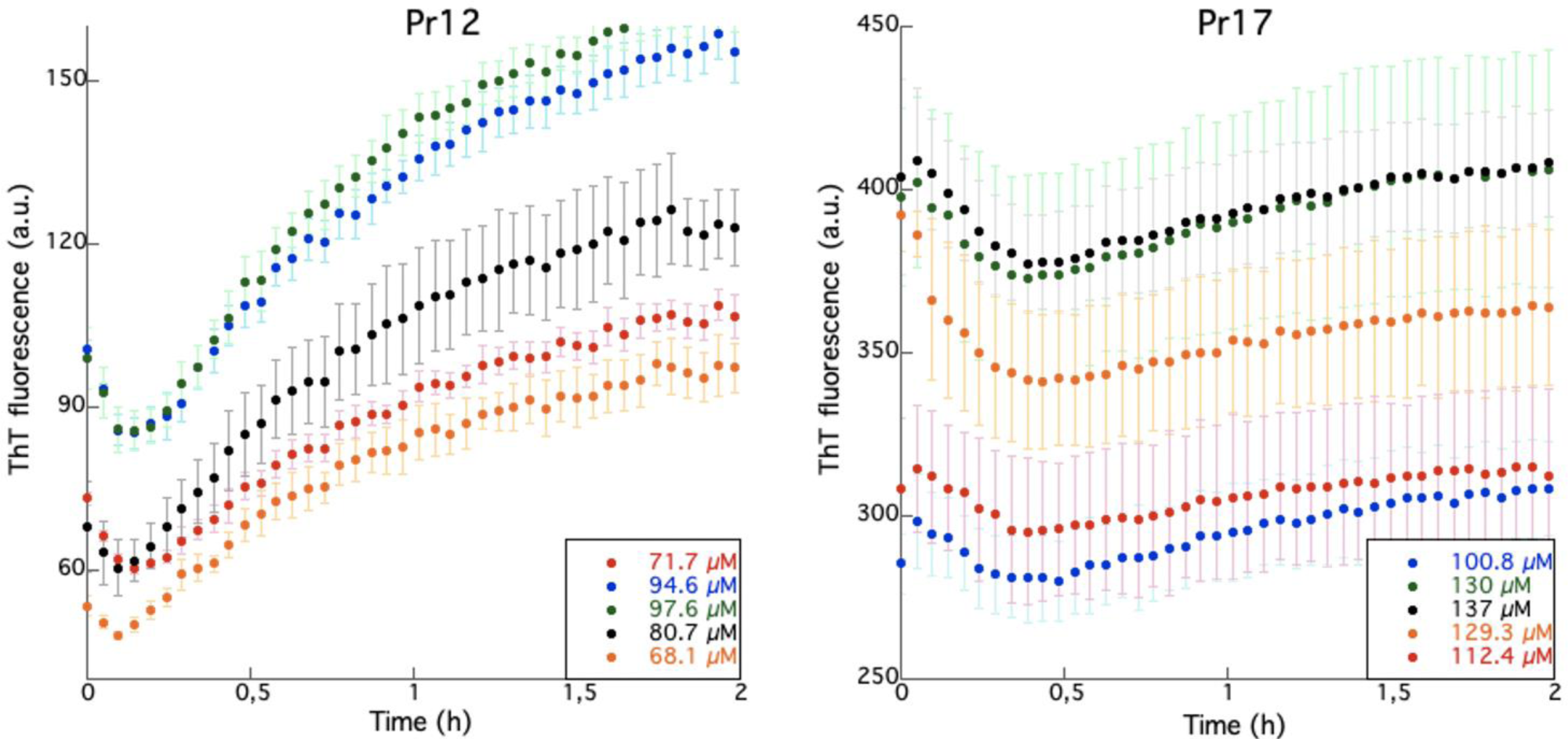
Blow up of the first 2 h of the signal change in Fig. 5. An initial dip in ThT fluorescence around t = 0 is observed for all samples.

**Figure S5.**
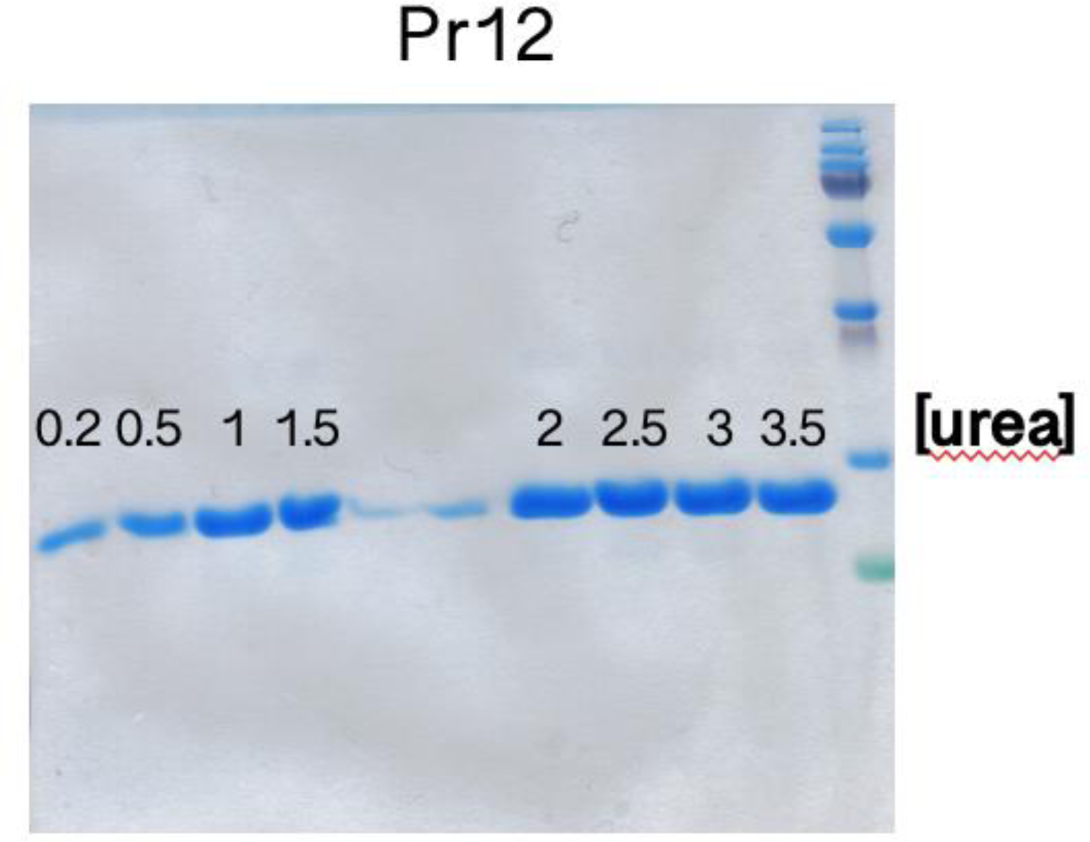
Supernatants of Pr12 samples after fibrillating 0.5 mg/mL protein in the presence of different concentrations of urea. The lanes between 1.5 M and 2 M urea are empty with some spill over from the other lanes.

**Figure S6.**
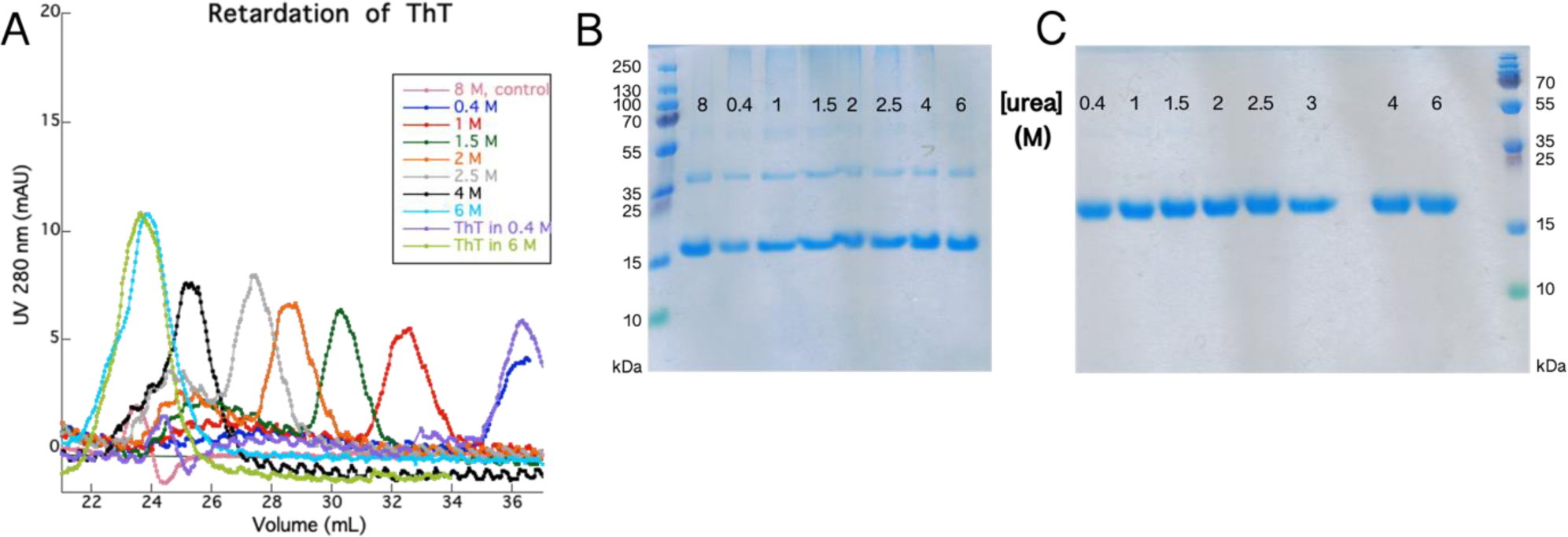
(**A**) Retardation of the ThT elution peak with increasing urea concentrations (and thereby increasing viscosity). Pr17 samples from the SEC experiments were analyzed with SDS-PAGE using either non-reducing (**B**) or reducing (**C**) loading buffer.

**Figure S7.**
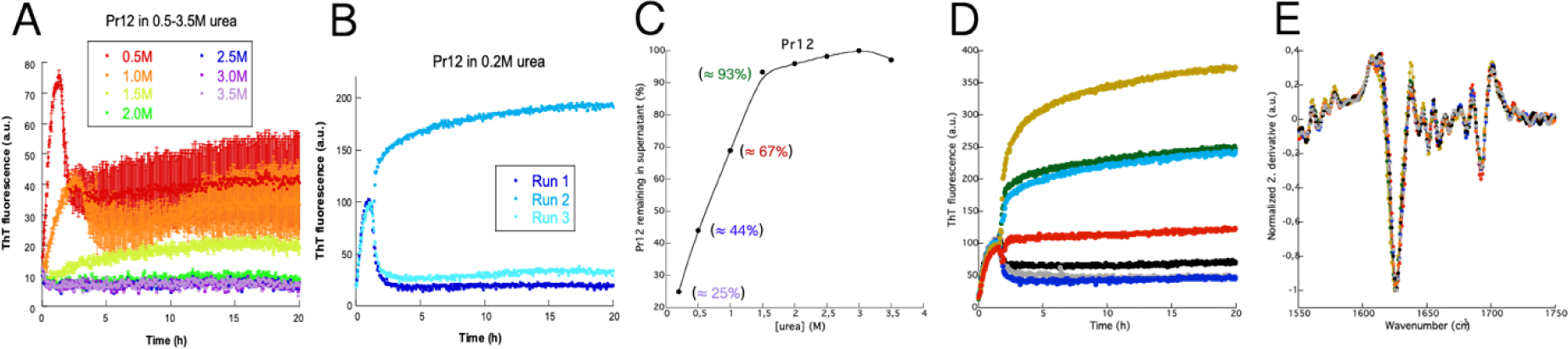
Aggregation of Pr12 at pH 7.4 in the presence of (A) 0.5-3.5 M urea or (B) 0.2 M urea (B) using a 60 mM Tris buffer instead of 1X PBS. (C) Amount of soluble protein left after aggregation was quantified by SDS-PAGE. (D) Seven samples containing the same solution of 0.5 mg/mL Pr12 were fibrillated to check if the differences in ThT end level was a consequence of different aggregate structures being formed. (D) The seven samples from panel D gave rise to identical FTIR spectra.

**Figure S8.**
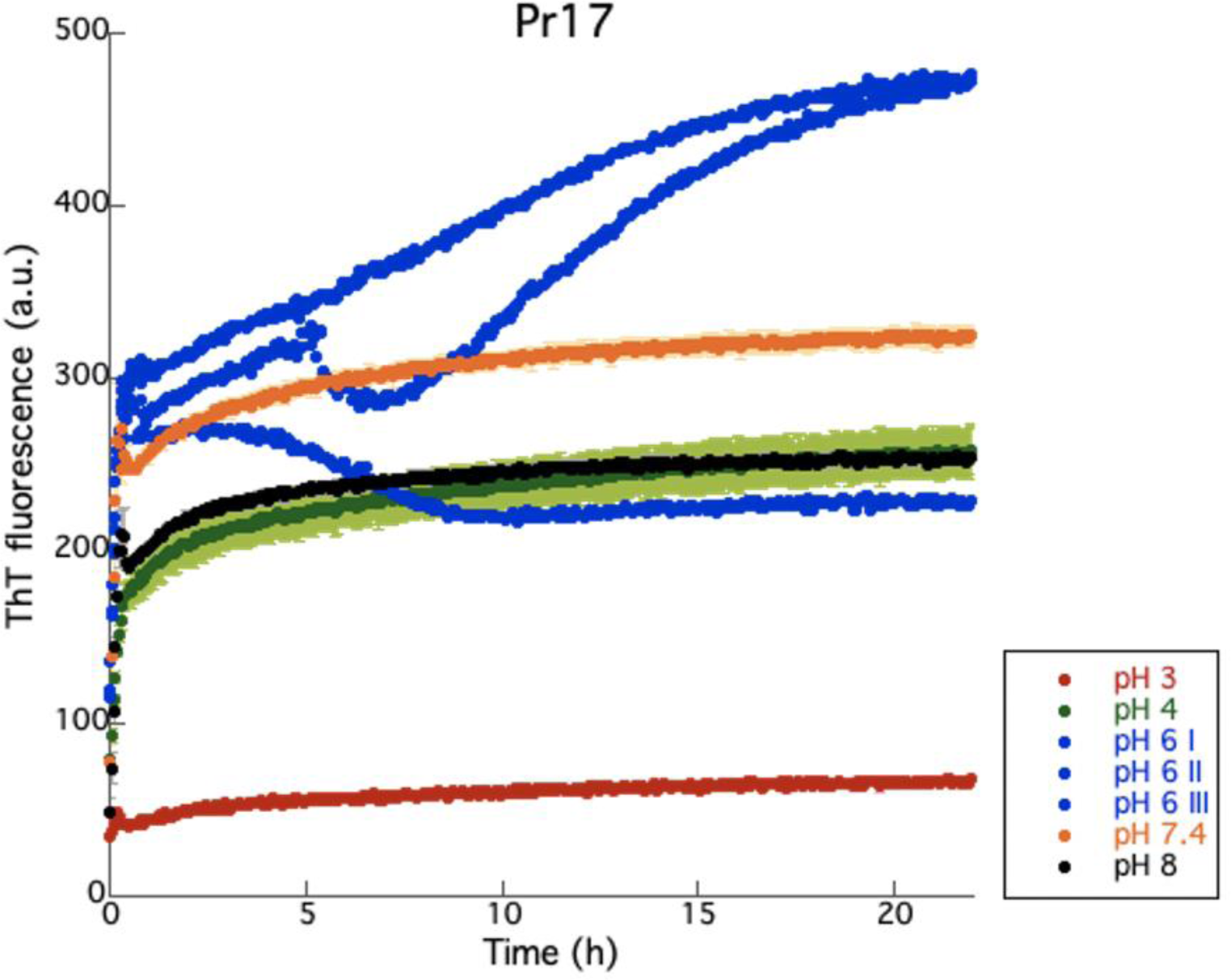
Aggregation of Pr17 at different pH values. Blue curves (pH 6) show highly irreproducible aggregation kinetics.

**Figure S9.**
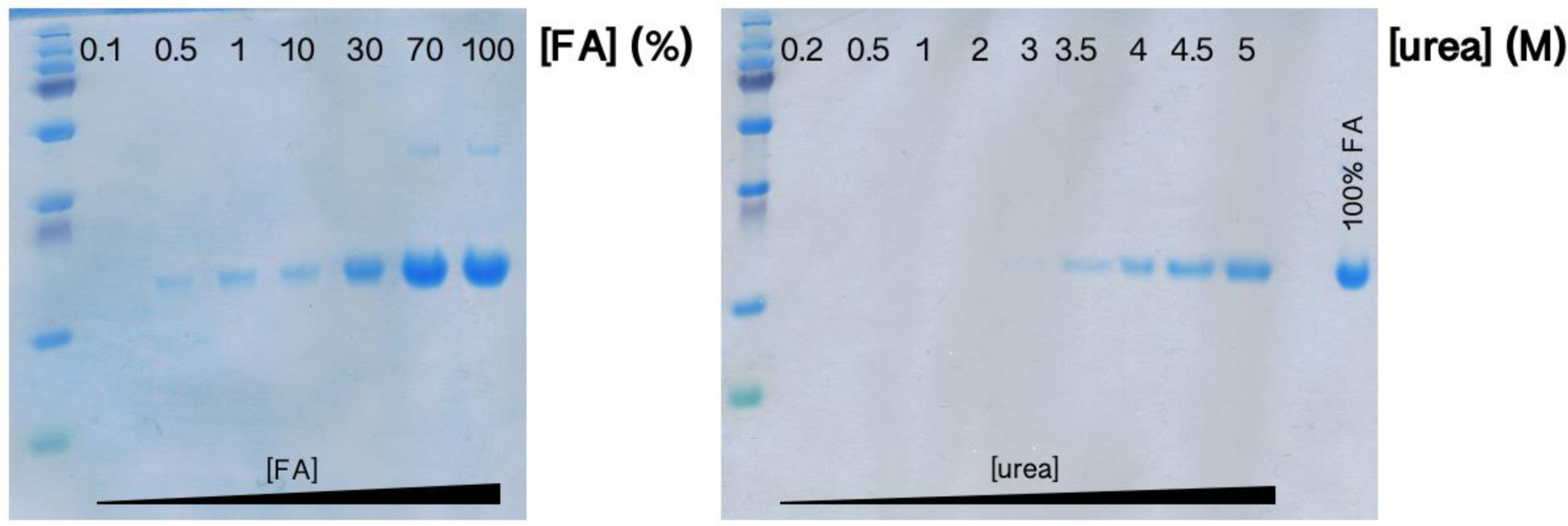
Dissolution of Pr17 aggregates by incubation with increasing concentrations of formic acid (left) or urea (right).

**Figure S10.**
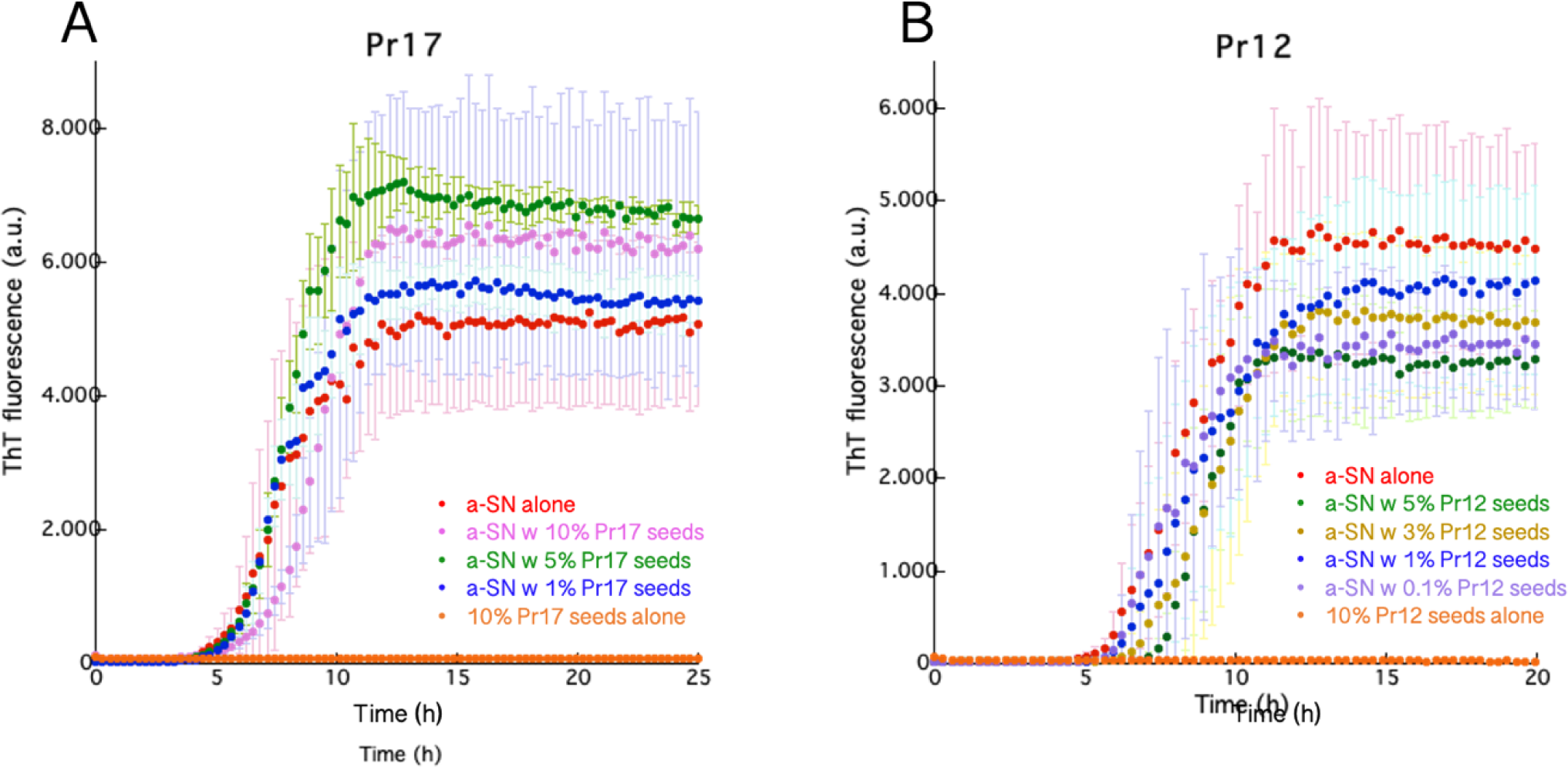
No cross-interaction is observed between monomeric αSN and seeds made of (**A**) Pr17 and (**B**) Pr12. 1 mg/mL αSN was fibrillated in the presence of different concentrations (percentage mass/mass) of Pr12/Pr17 seeds. The experiments were done in triplicates.

